# Renalase is a novel tissue and serological biomarker in pancreatic ductal adenocarcinoma

**DOI:** 10.1101/2021.04.12.439422

**Authors:** Yasheen Gao, Melinda Wang, Xiaojia Guo, Joanna Hu, Tian-min Chen, Sade’ M.B. Finn, Jill Lacy, John W. Kunstman, Charles H. Cha, Melena D. Bellin, Marie E. Robert, Gary V. Desir, Fred S. Gorelick

## Abstract

Dysregulated expression of the secretory protein renalase (RNLS) can promote pancreatic ductal adenocarcinoma (PDAC) growth in animal models. We characterized RNLS expression in premalignant and malignant PDAC tissue and investigated whether plasma RNLS levels corresponded to clinical PDAC characteristics. RNLS immunohistochemistry was used to determine the presence and distribution of RNLS in normal pancreas, chronic pancreatitis, PDAC precursor lesions, and PDAC tissues. Associations between pretreatment plasma RNLS and PDAC clinical status were assessed in patients with varied clinical stages of PDAC and included tumor characteristics, surgical resection in locally advanced/borderline resectable PDAC, and overall survival. Data were retrospectively obtained and correlated using non-parametric analysis. Mild to no RNLS was detected by histochemistry in the normal pancreas in the absence of abdominal trauma. In chronic pancreatitis, RNLS immunoreactivity localized to peri-acinar spindle-shaped cells in some samples. It was also widely present in PDAC precursor lesions and PDAC tissue. Among 240 patients with PDAC, elevated plasma RNLS levels were associated with worse tumor characteristics, including greater angiolymphatic invasion (80.0% vs. 58.1%, p = 0.012) and greater node positive disease (76.5% vs. 56.5%, p = 0.024). Overall survival was worse in patients with high plasma RNLS levels with median follow-up of 27.70 months vs. 65.03 months (p < 0.001). RNLS levels also predicted whether patients with locally advanced/borderline resectable (LA/BR) PDAC underwent resection (AUC 0.674; 95%CI 0.42-0.82, p = 0.04). Overall tissue RNLS was increased in both premalignant and malignant PDAC tissues compared to normal pancreas. Elevated plasma RNLS levels were associated with advanced tumor characteristics, decreased overall survival, and reduced resectability in patients with LA/BR PDAC. These studies show that RNLS levels are increased in premalignant pancreatic tissues and that its levels in plasma correspond to the clinical behavior of PDAC.

## Introduction

Pancreatic ductal adenocarcinoma (PDAC) is the seventh leading cause of cancer death worldwide with an estimated 430,000 annual deaths and an overall survival rate of less than 10%(1, 2). PDAC is often not detected until after it has disseminated systemically, is resistant to drug therapies, and is recognized as an urgent unmet medical need(3, 4). A better understanding of the biology and requirements for PDAC growth is crucial to improving the diagnostic and therapeutic approaches to this disease.

Renalase (RNLS) is a novel secretory protein (5-7) that can be highly expressed in PDAC tissue(8). Recombinant RNLS administration promotes the growth of human PDAC cell lines, and its inhibition leads to experimental tumor cell death in both in vitro and in vivo (8). Overall PDAC patient survival inversely correlated with tissue RNLS expression levels in tumor samples (8). These studies suggest that RNLS could be an important factor in maintaining PDAC growth and influence patient outcomes. Whether RNLS activity is associated with PDAC development, including in precursor lesions, has not been explored.

PDAC arises from several premalignant lesions including chronic pancreatitis (13), pancreatic intraepithelial neoplasia (PanIN), and pancreatic cystic neoplasms. PanIN (9), the most common predisposing lesion, progresses step-wise from non-invasive neoplasms (grades 1-3) to invasive PDAC with KRAS mutations being one of the earliest genetic events (10-12). Pancreatic cystic neoplasms, comprising intraductal papillary mucinous neoplasms (IPMNs) and less frequently, mucinous cystic neoplasms (MCNs), can also progress to PDAC (13).

Since RNLS is a secretory protein, changes in tissues levels could be reflected in its plasma levels. As seen with other tumors and biomarkers (e.g., CA19.9, CEA and PSA), it is possible that plasma RNLS levels could parallel tissue levels of RNLS in PDAC and serve as a new biomarker in PDAC.

Here we show that RNLS is present in premalignant (chronic pancreatitis, PanINs, and pancreatic cystic neoplasm) and malignant PDAC tissue suggesting a potential role for pancreatic RNLS in PDAC development. We also observed that elevations of a distinct form of plasma RNLS appear to correspond to PDAC prognosis in a subset of pancreas cancer patients with locally advanced disease at presentation.

## Methods

### Tissue specimens

Human pancreatic tissue samples from resection specimens of normal pancreas, chronic pancreatitis, neuroendocrine tumors, PDAC precursor lesions (PanIN, IPMN, and MCN), and PDAC, were obtained from Yale Surgical Pathology (Yale HIC: 2000021579). Additional chronic pancreatitis tissue samples were obtained from University of Minnesota Medical Center (IRB Approval Number: 0609M91887). Studies were given exceptions by the Veterans Administration HIC and Research committees under protocols FG0011-2020 and FSG0012-2020 (Oct 28, 2020).

### Immunohistochemistry Protocol

Immunohistochemistry was performed as described (8). Sections (5-μm) from formalin-fixed paraffin-embedded tissues were mounted on slides which were de-paraffinized and hydrated, followed by antigen retrieval in a pressure cooker containing citrate buffer (1g NaOH, 2.1 g citric acid in 1 L H2O pH 6). Sections were blocked with DAKO Dual endogenous enzyme block (Agilent, Santa Clara, CA, USA) for 10 min and 2.5% normal horse serum for 1 hr followed by incubation with primary antibody and isotype control IgG overnight at 4 °C. The m28-RNLS monoclonal antibody was diluted at 1µg/mL in buffer (TBS/1% Tween with 300 mM NaCl, pH = 7.4). ImmPRESS peroxidase-anti-rabbit IgG (Vector Laboratories, Burlingame, CA, USA) was used to detect primary antibodies. The color was developed using a Vector DAB substrate kit and tissue was counterstained with hematoxylin (Vector Laboratories). Hematoxylin and eosin stained and RNLS IHC stains were examined and at the light microscope to confirm the histologic diagnosis and document the distribution of RNLS immunopositivity by a pancreaticobiliary pathologist (MER). RNLS IHC staining was categorized as present or absent in all components of the pancreas (benign exocrine, endocrine, pancreatic ducts and stroma; neoplastic tissues, either in-situ or invasive). The specificity of labeling on each tissue was confirmed by performing labeling in tissue sections in the absence of the m28-RNLS primary antibody as well as using m28-RNLS antibody that had been pre-incubated with the peptide antigen (RP-220) (14).

### Immunofluorescence Protocol

The slides were deparaffinized and unmasked in the same manner as described in the immunohistochemistry protocol above. Following unmasking, sections were blocked with TBS/0.3% TritonX-100/10% goat serum) for 1 hr and incubated with a cocktail of m28-RNLS (1:100) plus α-smooth-muscle actin (αSMA) mouse monoclonal (1:400 Sigma-Aldrich, St. Louis, MI, USA) at 4°C overnight. Alexa 488-conjugated goat anti-rabbit (1:2000, Invitrogen Corporation, Carlsbad, CA) and Alexa 594-conjugated goat anti-mouse (1:2000, Invitrogen Corporation, Carlsbad, CA) were used to detect the primary antibodies. The sections were then incubated with an autofluorescence quenching kit (Vector Laboratories, Burlingame, CA, USA). The specificity of labeling on each tissue was confirmed by performing labeling in tissue sections in the absence of the m28-RNLS and αSMA primary antibodies.

### Measurement of plasma RNLS and CA-19.9 levels

Plasma RNLS levels were determined using the denaturation acid-sensitive pool and non-acid treated pool by ELISA as described (15). Here we refer to the acid-treated value as RNLS and the untreated value as non-acid treated RNLS. The median value was used to separate low and high RNLS levels. Serum CA19-9 levels were obtained from medical records before the first date of PDAC treatment.

Because of potentially spurious CA19-9 values in patients with biliary obstruction, patients with total bilirubin > 2.0 mg/dL were excluded from the CA19-9 analysis. CA19-9 levels of < 10 U/mL were also excluded from analysis because artifactually low CA19-9 levels can be seen in patients, especially those who have Lewis a-b-blood group antigen (16). A threshold CA19-9 value of 300 U/mL was designated elevated based on studies showing that pre-operative CA19-9 values above 300 U/mL suggest advanced disease and unresectable cancer (17).

### Clinical Data Collection

Prospectively collected plasma samples were obtained from consented patients with pathologically confirmed PDAC prior to initiation of treatment as part of the intake procedure for the Yale Gastrointestinal Tumor Biorepository from April 2012 to March 2019 (YGTB; Yale HIC #1203009817). Following definitive treatment, clinical and pathologic information were used to retrospectively annotate the plasma samples. Sociodemographic and clinicopathologic data were extracted from oncology visit notes, operation reports, and surgical pathology reports. Management was determined by the treating oncology team. Patients with locally advanced/borderline resectable (LA/BR) PDAC were determined at the time of diagnosis by cross-sectional imaging using a dedicated pancreatic contrast administration protocol and reviewed by a multidisciplinary PDAC management team that included an experienced pancreatic surgeon. Specifically, definitions of “resectable”, “borderline resectable”, and “locally advanced” were ascribed to patients by treating surgeons in accordance with the published National Comprehensive Cancer Network (NCCN) guidelines germane to the year of diagnosis (18). For the purposes of this study, patients who were identified as LA/BR at diagnosis who underwent resection with curative intent were considered “resectable” versus those who did not undergo surgery with curative intent. Any patients undergoing surgery for palliative purposes were not considered resectable.

### Statistical analyses

Nonparametric statistical analyses were performed using SPSS Version 24 (IBM Statistics; Armonck, NY). Bivariate nonparametric analyses of sociodemographic and treatment-related variables were performed using the calculated cut-off value for RNLS. Multivariate analyses were performed for statistically significant variables within the bivariate analysis. Sensitivity and specificity values were calculated using the calculated cut-off value for RNLS and pre-determined cut-off value for CA19-9. Kaplan-Meier survival analyses were performed using log rank analysis to calculate statistical significance to calculate overall survival and resectability of patients. All quantitative data were reported as median (range) or mean when median was unavailable and a p-value < 0.05 was used to determine statistical significance.

## Results

### RNLS immunoreactivity in human pancreatic tissue

We examined the presence and distribution of RNLS in normal human pancreas (n = 11), chronic pancreatitis (n = 32), PDAC precursor lesions (PanIN: n = 5, IPMN: n = 6, MCN: n = 4), and PDAC (n = 9) by immunohistochemistry. Sociodemographic and labeling characteristics for the patients examined are summarized in **Table 1a** and **Table 1b**.

**Table 1a.** Sociodemographic characteristics for RNLS immunohistochemistry patients.

| Characteristics | Histological variant |  |  |  |  |  |
| --- | --- | --- | --- | --- | --- | --- |
|  | Normal Pancreas (n = 11)<sup>a</sup> | Chronic Pancreatitis (n = 32)<sup>b</sup> | PanIN (n = 5) | IPMN (n = 6) | MCN (n = 4) | PDAC (n = 9) |
| Age<sup>c</sup>, years | 70 (23-94) | 56.8 (27-85) | 71 (62-82) | 70 (44-78) | 52 (26-74) | 67 (51-80) |
| Male<sup>d</sup> | 4 (36.4) | 13 (40.6) | 2 (40.0) | 4 (66.7) | 0 | 5 (55.6) |
| White<sup>d</sup> | 7 (63.6) | 28 (87.6) | 5 (100) | 5 (83.3) | 4 (80) | 8 (88.9) |
| BMI<sup>c</sup>, kg/m<sup>2</sup> | 28.0 (25.8-30.1) | 24.3 (13.3-37.1) | 26.0 (18.9-28.7) | 28.4 (14.9-39.6) | 22.6 (13.3-34.1) | 30.4 (23.3-34.9) |
| Current or past smoker<sup>d</sup> | 4 (36.4) | 17 (53.1) | 3 (60) | 4 (66.7) | 2 (40) | 5 (55.6) |
| Alcohol use<sup>d</sup> | 0 | 5 (15.6) | 0 | 0 | 1 (20) | 1 (11.1) |
| Hypertension<sup>d</sup> | 3 (27.3) | 13 (40.6) | 2 (40) | 2 (33.3) | 1 (20) | 3 (33.3) |
| Cardiovascular Disease<sup>d</sup> | 0 | 3 (8.4) | 1 (20) | 0 | 0 | 1 (11.1) |
| Chronic Kidney Disease<sup>d</sup> | 0 | 2 (6.3) | 0 | 0 | 1 (20) | 1 (11.1) |
| Diabetes<sup>d</sup> | 0 | 9 (28.1) | 3 (60) | 2 (33.3) | 2 (40) | 2 (22.2) |

<sup>a</sup> Seven normal pancreas cases were taken from patients who had undergone Whipple surgery for duodenal adenoma. Four normal pancreas cases were taken from patients who had undergone splenectomy.
<sup>b</sup> Five cases were taken from patients who had alcohol induced chronic pancreatitis. Four cases were taken from patients with idiopathic chronic pancreatitis. Six cases were taken from patients who had genetic chronic pancreatitis. One case came from a patient who had chronic pancreatitis due to pancreatic divisum. One case was taken from a patient who had chronic pancreatitis due to Sphincter of Oddi dysfunction. Two cases were taken from patients who had chronic pancreatitis associated with pseudocyst. Thirteen cases were taken from patients with PDAC precursor lesions and PDAC.
<sup>c</sup> Data are given as median (range)
<sup>d</sup> Data are given as number (percentage)

**Table 1b.** Labeling characteristics for RNLS immunohistochemistry patients.

| Pancreatic tissue type | n | Benign ductal epithelium | Acinar cell | Islets | Neoplastic cells | Stellate cells |
| --- | --- | --- | --- | --- | --- | --- |
| Normal pancreas (trauma) | 4 | 4/4 (100) | 4/4 (100) | 4/4 (100) | N/A | 1/4 (25) |
| Normal pancreas (duodenal adenoma) | 7 | 1/7 (14) | 3/7 (43) | 1/7 (14) | N/A | 0/7 (0) |
| Chronic pancreatitis (Non-tumor associated) | 19 | 6/19 (32) | 8/19 (42) | 6/19 (32) | N/A | 6/19 (32) |
| Chronic pancreatitis (adjacent to pancreatic precursor lesion/PDAC) | 13 | 12/13 (92) | 13/13 (100) | 12/13 (92) | N/A | 2/13 (15) |
| Precursor Lesion (PanIn) | 5 | 5/5 (100) | 5/5 (100) | 5/5 (100) | 5/5 (100) | 0/5 (0) |
| Precursor Lesion (IPMN) | 6 | 6/6 (100) | 6/6 (100) | 5/6 (83.3) | 6/6 (100) | 0/6 (0) |
| Precursor Lesion (MCN) | 4 | 4/4 (100) | 4/4 (100) | 4/4 (100) | 4/4 (100) | 0/4 (0) |
| PDAC | 9 | 4/9 (44) | 4/9 (44) | 5/9 (55) | 9/9 (100) | 0/9 (11) |

Among normal human pancreatic tissue from patients who had undergone Whipple procedure for duodenal adenoma (n = 7), there was little to no RNLS labeling (**Figure 1a**). In patients who had undergone distal pancreatectomy associated with trauma-related splenectomy (n = 4) there was granular RNLS labeling in the apical cytoplasm of pancreatic acinar cells as well as diffuse RNLS labeling in pancreatic islets and ducts (**Figure 1b)**. One of these tissues showed RNLS labeling in spindle-shaped cells surrounding pancreatic acinar cells, a cellular distribution more often seen in chronic pancreatitis.

**Figure 1.**
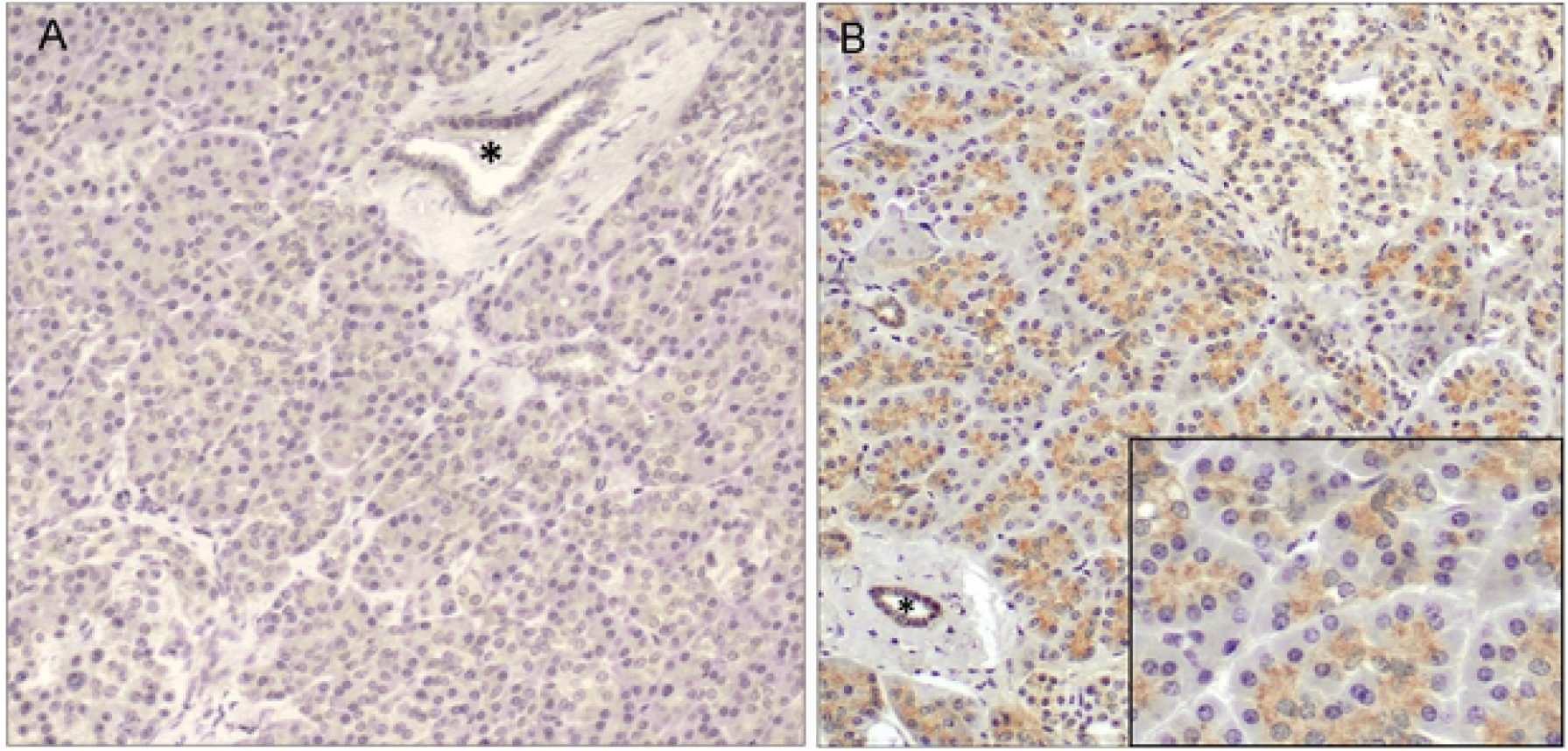
RNLS labeling in normal human pancreas. Representative images of RNLS labeling detected by immunohistochemistry using the m28-RNLS antibody in the normal human pancreas from (A) a patient who had undergone Whipple procedure for duodenal adenoma with essentially no signal and (B) a patient who had undergone splenectomy for trauma, showing granular cytoplasmic staining. The asterisk indicates a normal pancreatic duct. Original magnification 200x; inset 400x

Chronic pancreatitis tissues were obtained from patients with the following etiologies: alcoholic chronic pancreatitis (n = 5), genetic chronic pancreatitis (n = 6), idiopathic chronic pancreatitis (n = 4), chronic pancreatitis due to pancreatic divisum (n = 1), chronic pancreatitis due to Sphincter of Oddi dysfunction (n = 1), chronic pancreatitis adjacent to a benign inflammatory pseudocyst (n = 2), chronic pancreatitis associated with precursor lesions (n = 9), and chronic pancreatitis associated with PDAC (n = 4).

Overall, RNLS labeling in chronic pancreatitis was present, but variable, as noted below and in **Table 1b**. Using immunocytochemistry, we found RNLS labeling in spindle-shaped cells surrounding pancreatic acinar cells in six of the nineteen non-tumor associated chronic pancreatitis cases (**Figure 2a**) and two of the thirteen tumor-associated chronic pancreatitis cases (**Figure 2b**). These cases included one case each of the following etiologies: 1) Mutations in the cystic fibrosis transmembrane conductance regulator (CTFR), 2) Mutation in chymotrypsin C (CTRC), 3) Idiopathic chronic pancreatitis, 4) Alcoholic chronic pancreatitis, 5) Pancreatic divisum, 6) Associated with a pseudocyst, 7) Associated with MCN and 8) Chronic pancreatitis associated with PDAC. To determine the identity of the spindle-shaped containing RNLS immunoreactivity, we performed double-label (RNLS and αSMA-smooth muscle actin) immunofluorescence on six non-tumor associated and three tumor associated chronic pancreatitis cases In a chronic pancreatitis case (associated CTRC mutation) we observed co-localization of RNLS and αSMA (**Figure 2c-e**), suggesting that RNLS may be present in stellate cells. In the remainder of the cases the labeling was faint, and it was unclear whether the markers co-distributed. In non-tumor associated chronic pancreatitis samples RNLS immunoreactivity was also observed in acinar cells (8/19 samples), pancreatic ducts (6/19) and islet cells (6/19). (**Table 1b**). Granular cytoplasmic RNLS labeling of acinar cells, pancreatic ducts and islet cells was also present in regions of chronic pancreatitis associated with neoplastic conditions, including PanIN (n = 2), IPMN (n = 4), MCN (n = 3), and PDAC (n = 5). Finally, RNLS labeling was noted in stromal cells, including mononuclear inflammatory cells and fibroblasts, in regions of chronic pancreatitis associated with neoplastic precursor lesions in four samples.

**Figure 2.**
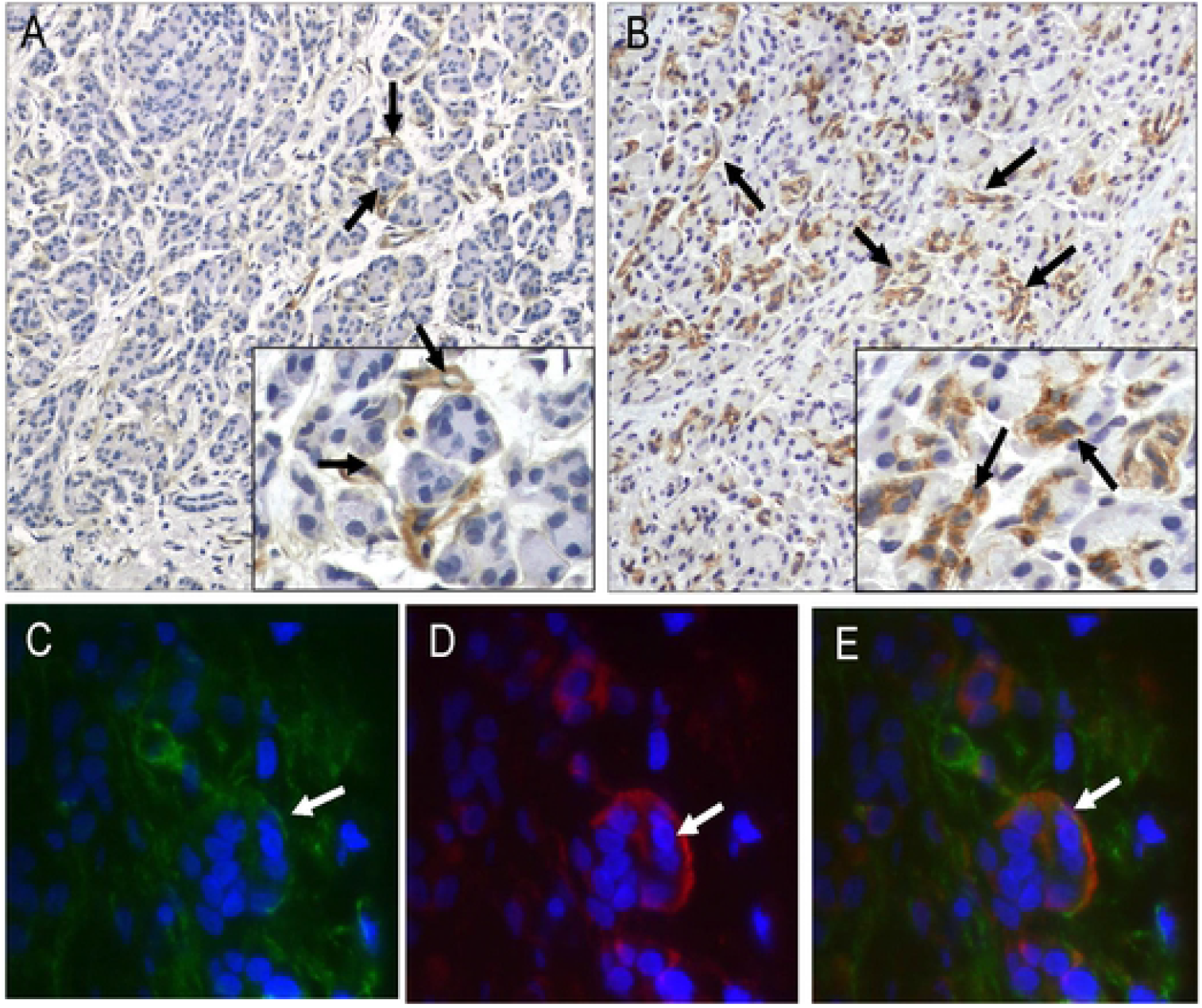
RNLS is present in spindle-shaped cells in human chronic pancreatitis. Representative images of RNLS labeling detected by immunohistochemistry using the m28-RNLS antibody in (A) genetic chronic pancreatitis (CTRC mutation) and (B) PDAC-associated chronic pancreatitis. (C-E) Immunofluorescence labeling of RNLS with m28-RNLS antibody (green) and alpha-smooth muscle actin (red) in a patient with genetic chronic pancreatitis (CTRC mutation). The white arrows point to cells positive for both proteins (stellate cells). Original magnification A and B: 200x, insets 400x. Original magnification C-E: 10000x.

We labeled for RNLS immunoreactivity in PDAC precursor lesions (n = 15) including PanIN (n = 5), IPMN (n = 6), and MCN (n = 4), and in PDAC tumors (n = 9). In all premalignant and malignant PDAC tissues there was diffuse RNLS labeling in in-situ and invasive neoplastic epithelium, adjacent benign duct cells, acinar cells, and islets (**Figure 3a-d**). In six of the fifteen precursor lesion cases and three of the nine PDAC cases, RNLS labeling was noted in stromal cells (comprising fibroblasts, endothelial cells and inflammatory cells). (**Figure 3e**) In-situ and invasive pancreatic ductal neoplasms and precursor lesions showed increased RNLS staining compared to benign samples (normal pancreas and most cases of pancreatitis). Specifically, in precursor and PDAC samples, RNLS was present in the majority of cell types, including all benign elements. The intensity of labeling often appeared more intense at light microscopy in neoplastic than in benign epithelium but was not quantified because of limitations of the labeling technique.

**Figure 3.**
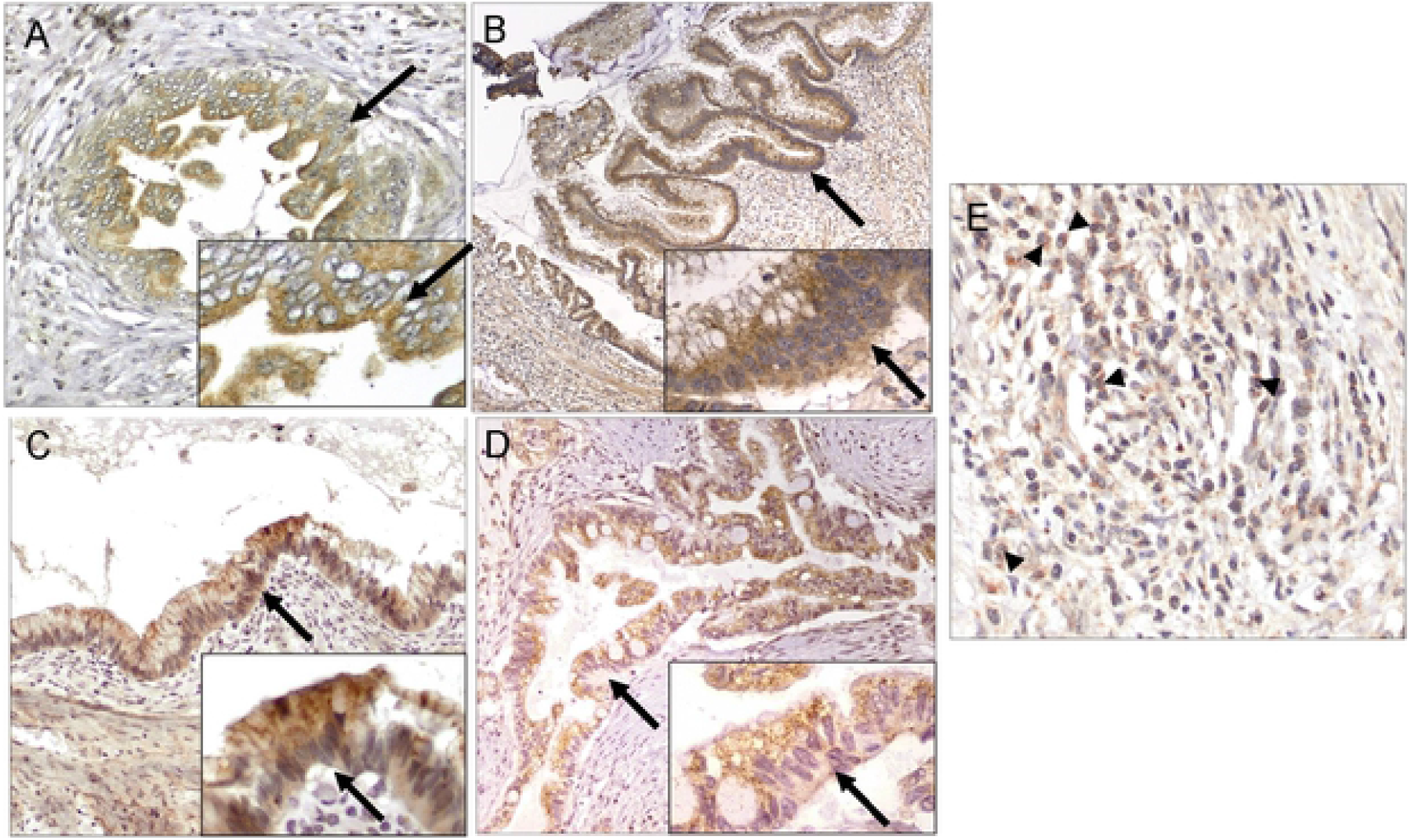
RNLS distribution in human precursor and PDAC tissue. Representative images of RNLS labeling detected by immunohistochemistry using the m28-RNLS antibody in: (A) PanIN, (B) IPMN, (C) MCN, and (D-E) PDAC. Intense granular cytoplasmic staining is present in the majority neoplastic epithelial cells, and in a subset of mononuclear inflammation near tumor. The black arrows point to ductal cells in (A) PanIN, (B) IPMN, (C) MCN, and (D) PDAC. The black arrowheads point to mononuclear inflammatory cells in the PDAC stroma. Original magnification: A, E 400x, inset 600x; B 100x, inset 600x; C,D 200x, insets 400x.

### Plasma RNLS in patients with PDAC

Three hundred and forty-seven patients with biopsy confirmed PDAC were identified. Of those, 240 patients had both plasma RNLS levels drawn prior to any PDAC treatment and medical records available for review. Median RNLS level was 1329.15 ng/ml. High plasma RNLS was associated with younger age at diagnosis but not associated with race, gender, or BMI. Patients with higher serum protein levels at diagnosis also had higher plasma RNLS levels at diagnosis (7.30 vs. 7.00, p = 0.004). Patients with high RNLS levels also exhibited worse PDAC disease attributes (**Table 2**). This included higher angiolymphatic invasion (80.0% vs. 58.1%, p = 0.012), and greater node positive disease (76.5% vs. 56.5%, p = 0.024) particularly hepatic artery lymph node positivity (26.7% vs. 6.7%, p = 0.022). Patients with higher plasma RNLS levels did not exhibit larger tumor size, or greater rates of margin positivity (**Tables 2**). In concordance with worse disease features, patients with higher plasma RNLS levels were less likely to be deemed clinically resectable at diagnosis (39.2% vs. 58.0%, p = 0.004) and were more likely to have disseminated disease (distant metastasis) at diagnosis (25.8% vs. 13.4%, p = 0.022). Higher plasma RNLS levels at diagnosis were associated with decreased survival with a median of 27.70 months vs. 65.03 months (p < 0.001; **Figure 4a**). The survival difference persisted when RNLS levels were divided into tertiles (n = 80 per group; Bottom tertile = 19.13 months, Middle tertile = 38.47 months, Top tertile = 65.03 months; p < 0.001; **Figure 4b**) and into quartiles (n = 60 per group; Bottom quartile = 19.13 months, Bottom middle quartile = 38.47 months, Top middle quartile = 70.43 months, Top quartile = 65.03 months, p < 0.001; **Figure 4c**). There was no correlation between non-acid treated RNLS levels and either tumor characteristics or survival (data not shown).

**Table 2.** Sociodemographic and clinicopathologic features among patients with PDAC with high vs. low plasma RNLS.

| *Sociodemographic Characteristics* |  |  |  |  |
| --- | --- | --- | --- | --- |
| Age at Diagnosis | 68.00 (9.83) | 66.00 (9.62) | 68.00 (9.80) | 0.018 |
| Race |  |  |  | 0.236 |
| White | 114/120 (95.0%) | 107/120 (89.2%) | 221/240 (92.1%) |  |
| Black | 5/120 (4.2%) | 10/120 (8.3%) | 15/240 (6.3%) |  |
| Other | 1/120 (0.8%) | 3/120 (2.5%) | 4/240 (1.7%) |  |
| Male | 71/120 (59.2%) | 75/120 (62.5%) | 146/240 (60.8%) | 0.692 |
| BMI at Diagnosis | 26.37 (5.29) | 26.34 (6.92) | 26.29 (6.04) | 0.719 |
| Albumin | 4.00 (0.50) | 4.00 (0.50) | 4.00 (0.49) | 0.237 |
| Total Protein | 7.00 (0.71) | 7.30 (0.69) | 7.10 (0.73) | **0.004** |
| *Clinicopathologic Characteristics* |  |  |  |  |
| Perineural Invasion | 61/75 (81.3%) | 46/50 (92.0%) | 107/125 (85.6%) | 0.122 |
| Angiolymphatic Invasion | 43/74 (58.1%) | 40/50 (80.0%) | 83/124 (66.9%) | **0.012** |
| Differentiation |  |  |  | 0.734 |
| Well | 3/75 (4.0%) | 1/50 (2.0%) | 4/125 (3.2%) |  |
| Moderate | 37/75 (49.3%) | 23/50 (46.0%) | 50/125 (48.0%) |  |
| Poor | 35/75 (46.7%) | 26/50 (52.0%) | 61/125 (48.8%) |  |
| Node Positive | 43/76 (56.6%) | 39/51 (76.5%) | 82/127 (64.6%) | **0.024** |
| HALN Positive | 3/45 (6.7%) | 8/30 (26.7%) | 11/75 (14.7%) | **0.022** |
| Tumor Size ≥ 2cm | 58/73 (79.5%) | 44/49 (89.8%) | 102/122 (83.6%) | 0.144 |
| Any Margin | 15 (19.7) | 15 (29.4) | 30 (23.6) | 0.237 |
| *Whipple Characteristics* | **Low (n=53)** | **High (n=36)** | **Total (n=89)** |  |
| Neck Margin | 1 (1.9) | 2 (5.6) | 3 (3.4) | 0.563 |
| Bile Duct Margin | 0 | 0 | 0 | -- |
| Anterior Circumferential Margin | 0 | 0 | 0 | -- |
| Posterior Circumferential Margin | 11 (20.8) | 6 (16.7) | 17 (19.1) | **0.785** |
| Vascular Groove | 2 (3.8) | 0 (0.0) | 2 (2.2) | **0.513** |
| Uncinate Process | 10 (18.9) | 6 (16.7) | 16 (18.0) | 1.000 |
| Duodenal | 0 (0.0) | 1 (2.8) | 1 (1.1) | 0.404 |
| Invades Duodenum | 25 (47.2) | 19 (52.8) | 44 (49.4) | 0.669 |
| Gastric | 0 | 0 | 0 | -- |
| Pancreatic Duct | 13 (24.5) | 13 (36.1) | 26 (29.2) | 0.248 |
| CBD | 12 (22.6) | 9 (25.7) | 21 (23.9) | 0.801 |
| Peripancreatic soft tissue | 43 (81.1) | 32 (88.9) | 75 (84.3) | 0.386 |
| Any margin | 12 (22.6) | 9 (25.0) | 21 (23.6) | 0.805 |
| Whipple | 53 (68.8) | 36 (70.6) | 89 (69.5) | 1.000 |
| *Distal Pancreatectomy Characteristics* | **Low (n=24)** | **High (n=15)** | **Total (n=39)** |  |
| Peripancreatic soft tissue | 16 (69.6) | 13 (86.7) | 29 (76.3) | 0.273 |
| Pancreatic | 0 | 0 | 0 | -- |
| Anterior | 1 (4.3) | 1 (6.7) | 2 (5.3) | 1.000 |
| Posterior | 2 (8.7) | 4 (26.7) | 6 (15.8) | 0.188 |
| Spleen | 0 (0.0) | 1 (6.7) | 1 (2.6) | 0.395 |
| Any Margin | 3 (13.0) | 6 (40.0) | 9 (23.7) | 0.115 |

**Figure 4.**
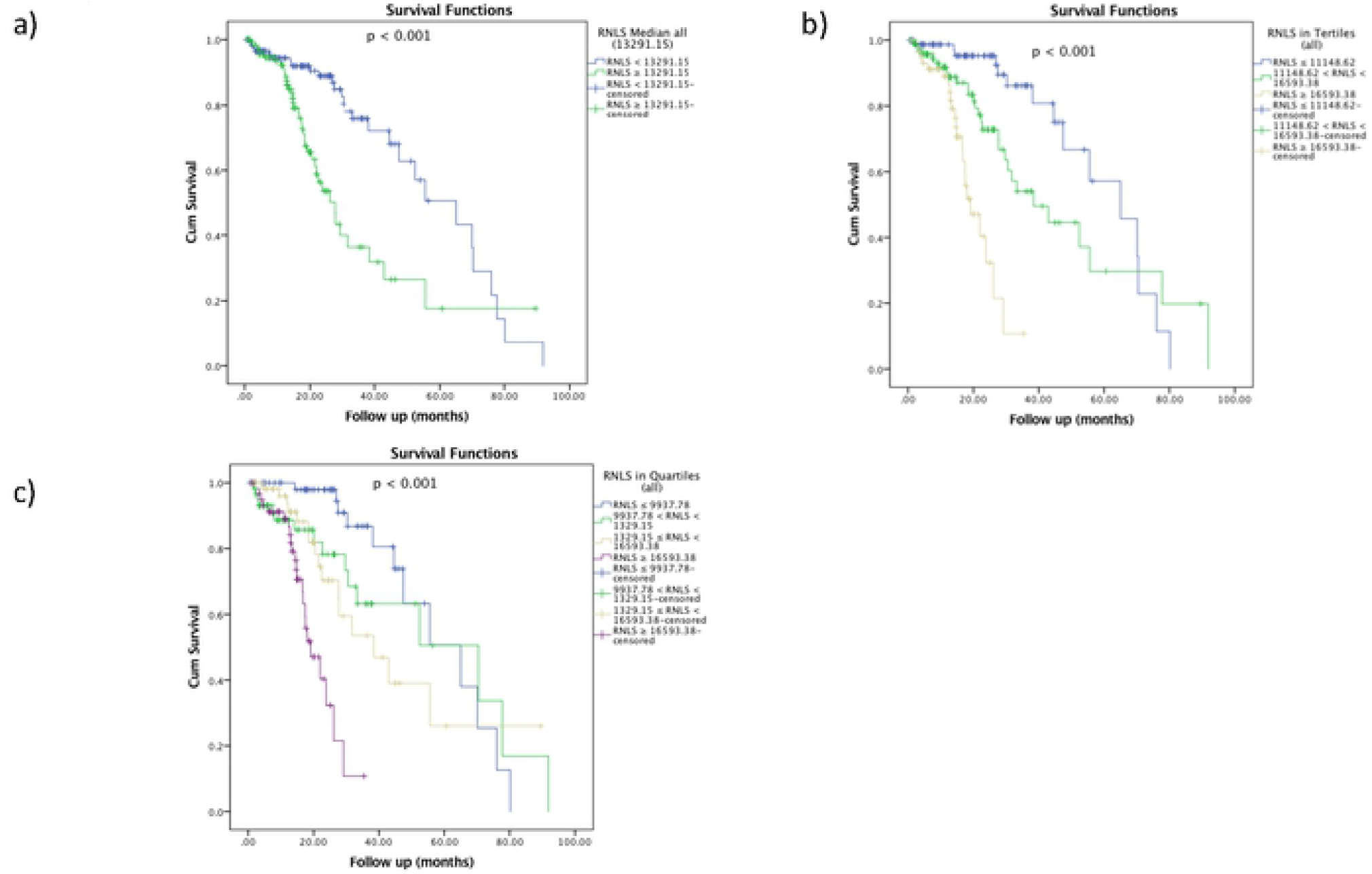
High plasma RNLS levels are associated with worse survival in patients with PDAC. (A) When RNLS is cut by median, (B) by tertiles, and (C) by quartiles

### Plasma RNLS in patients with locally advanced/borderline resectable PDAC

Among patients with PDAC and available plasma RNLS levels, 76 patients presented with LA/BR PDAC according to the treating oncology team at diagnosis. Median RNLS level among all LA/BR PDAC patients was 13601.55 ng/ml. Plasma RNLS levels did not correlate with age, race, gender, BMI, serum albumin, or total blood protein at diagnosis (**Table 3**). High RNLS level, defined as values above the median, did not correlate with high CA19-9 levels (**Table 3**). Of note, only 22 of the total 76 patients had CA19-9 values at diagnosis that met the exclusion criteria. Additionally, RNLS levels were not associated with the reported tumor size, tumor location, or vessel encasement on imaging at initial presentation (**Table 3**). Among LA/BR patients, high RNLS levels were associated with worse overall survival than were low RNLS levels (p = 0.015, **Figure 5a**). At a median follow-up of 19.60 months, median overall survival was 22.83 months among patients with low RNLS and 16.93 months among those with high RNLS levels. Survival correlations were especially discriminatory when RNLS levels were analyzed using tertiles (n = 25 per group; Bottom tertile = 70.13 months, Middle tertile = 27.70 months, Top tertile = 23.83 months; p = 0.004, **Figure 5b**) and quartiles (n = 19 per group; Bottom quartile = 60.63 months, Bottom middle quartile = 43.67 months, Top middle quartile = 25.81 months, Top quartile = 18.62 months; p < 0.001, **Figure 5c**). As a continuous variable, RNLS level was also associated with survival (p = 0.002). CA19-9 levels were not as predictive of survival. Median survival among those with low vs. high CA19-9 levels was 21.47 vs. 19.60 months (p = 0.548). There was no significant difference for any of these parameters when compared to values for non-acid treated RNLS (data not shown).

**Table 3.** Patient characteristics for low vs. high plasma RNLS levels in patients with locally advanced/borderline resectable (LA/BR) pancreatic adenocarcinoma (PDAC)

| Factor | Low RNLS (n=38) | High RNLS (n=38) | p-value |
| --- | --- | --- | --- |
| Age at Diagnosis | 67.0 (46-90) | 63.5 (37-86) | 0.651 |
| Race |  |  | 0.744 |
| White | 35 (92.1%) | 33 (86.8%) |  |
| Black | 2 (5.3%) | 3 (7.9%) |  |
| Other | 1 (2.6%) | 2 (5.3%) |  |
| Gender |  |  | 0.818 |
| Male | 20 (52.6%) | 22 (57.9%) |  |
| Female | 18 (47.4%) | 16 (42.1%) |  |
| Chronic Kidney Disease | 1 (2.6%) | 2 (5.3%) | 1.000 |
| Hypertension | 12 (31.6%) | 14 (36.8%) | 0.809 |
| Heart Failure | 1 (2.6%) | 0 (0.0%) | 1.000 |
| Coronary Artery Disease | 4 (10.5%) | 2 (5.3%) | 0.674 |
| BMI at Diagnosis | 25.45 (17.81-40.04) | 26.34 (16.46-68.11) | 0.875 |
| Albumin at Diagnosis | 4.10 (2.50-4.70) | 4.00 (3.20-4.70) | 0.232 |
| Total Protein at Diagnosis | 7.10 (5.60-8.20) | 7.15 (5.70-8.30) | 0.555 |
| High CA19-9 at Diagnosis <sup>a</sup> | 6 (50.0%) | 2 (20.0%) | 0.204 |
| Large Vessel Encasement | 10 (26.3%) | 12 (31.6%) | 0.801 |
| PDAC Location |  |  | 0.377 |
| Head | 31 (81.6%) | 28 (73.7%) |  |
| Body | 6 (15.8%) | 6 (15.8%) |  |
| Tail | 1 (2.6%) | 4 (10.5%) |  |
| PDAC Size by imaging | 30.50 (15.00-90.00) | 31.50 (18.00-92.00) | 0.705 |

<sup>a</sup> Only includes patients with total bilirubin < 2.0 and CA19-9 >10 (n = 22)

**Figure 5.**
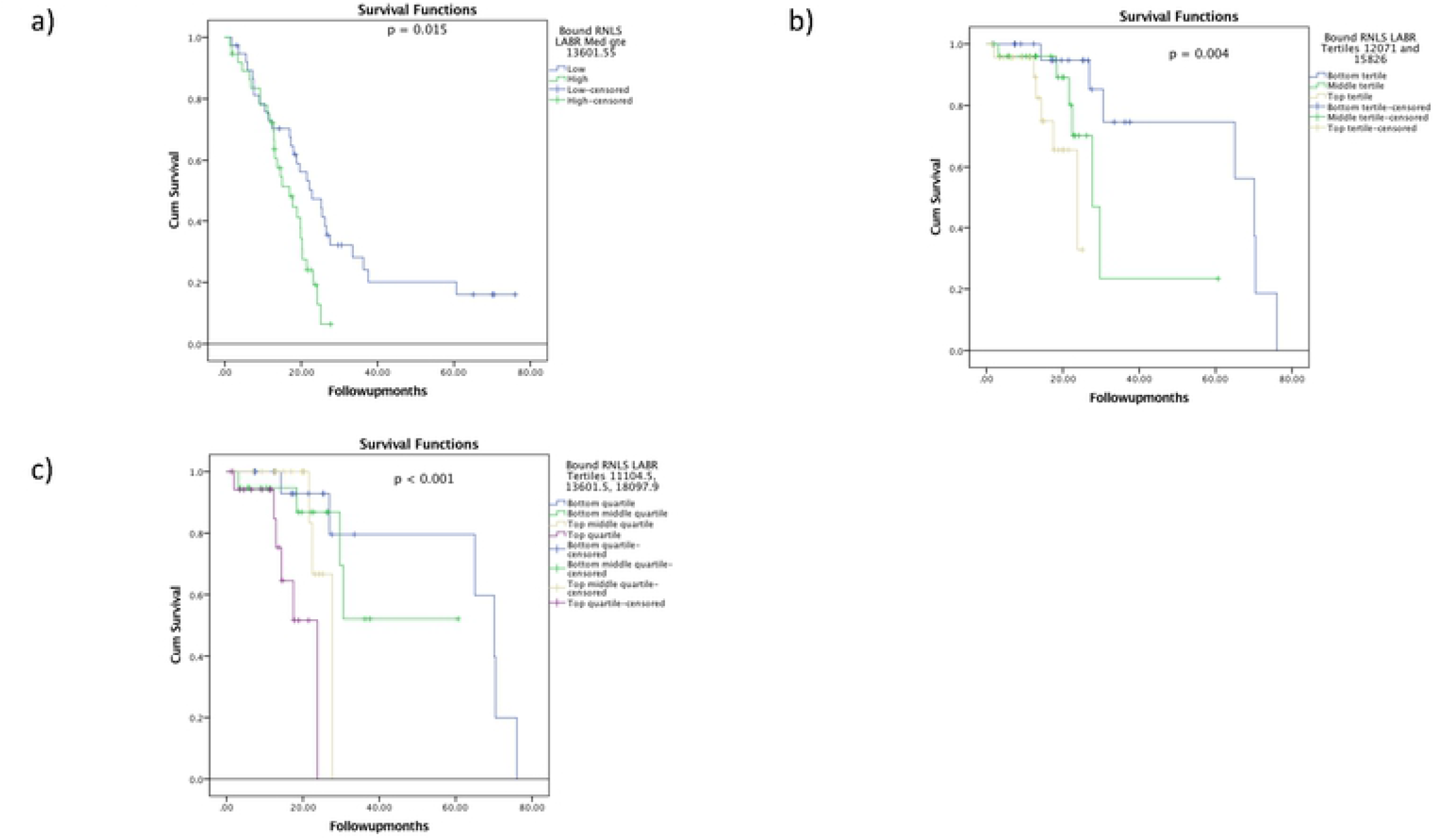
High plasma RNLS levels are associated with worse survival in patients with LA/BR PDAC. (A) When RNLS is cut by median, (B) by tertiles, and (C) by quartiles

Of the 76 patients with LA/BR PDAC, 18 (23.7%) underwent successful resection. Reasons for no eventual surgery in 58 patients were persistent local disease in 37 (48.7%), metastatic disease in 7 (9.2%), poor surgical candidacy in 4 (5.3%), and death in 3 (3.9%). The remaining 7 (9.2%) underwent exploratory laparoscopy and were subsequently found to have either unresectable local (5) or metastatic (2) disease upon surgical examination. Low RNLS levels were associated with greater odds of undergoing resection (aOR = 0.29 (0.09-0.93, p = 0.036, **Table 4**). Of patients with low RNLS levels, 34.2% underwent resection compared to 15.2% patients with high RNLS levels, a > 2-fold difference. On the other hand, serum CA19-9 levels showed no relationship to resection status (aOR = 0.33, p = 0.272, **Table 4**). RNLS levels among patients who underwent resection, versus no resection are shown in Figure 3.

**Table 4.** Treatments for high vs. low RNLS levels in patients with locally advanced/borderline resectable pancreatic adenocarcinoma as compared to CA19-9.

| **RNLS** |  |  |  |  |
| --- | --- | --- | --- | --- |
| **Factor** | **Low (n = 38)** | **High (n = 38)** | **aOR (95%CI)** | **p-value** |
| Resection | 13 (34.2%) | 5 (13.2%) | 0.29 (0.09-0.93) | **0.036** |
| Chemotherapy | 36 (94.7%) | 36 (97.3%) | 2.00 (0.17-23.06) | 0.578 |
| Radiotherapy | 18 (48.6%) | 21 (58.3%) | 1.48 (0.59-3.73) | 0.408 |
| **CA19-9** |  |  |  |  |
| **Factor** | **Low (n = 14)** | **High (n = 8)** | **aOR (95%CI)** | **p-value** |
| Resection | 3 (42.9%) | 3 (20.0%) | 0.33 (0.05-2.37) | 0.272 |
| Chemotherapy | 6 (85.7%) | 15 (100.0%) | -- | 1.000 |
| Radiotherapy | 5 (71.4%) | 7 (46.7%) | 0.35 (0.05-2.41) | 0.286 |

Further, ROC analysis shows that lower plasma RNLS levels at diagnosis predicted resection among patients with LA/BR disease (AUC (area under curve) = 0.674; 95% CI 0.53-0.82; p = 0.037, **Figure 6**. The cut-off value with the highest sensitivity and specificity as determined by ROC analysis, 12548.46 ng/mL, was similar to the median, 13601.55 ng/mL used as the cut-off for the designation of low vs high RNLS status. When using a cut-off value specified by AUC analysis, low RNLS was associated with resection with a sensitivity of 70.69% and a specificity of 66.67% (**Table 5**). Low CA19-9 levels showed lower sensitivity and lower specificity when compared to RNLS levels at AUC or median cut-off.

**Table 5.** Sensitivity and specificity tests of CA19-9 and RNLS levels to predict resection in patients with locally advanced/borderline resectable pancreatic adenocarcinoma.

|  | **Sensitivity (95% CI)** | **Specificity (95% CI)** |
| --- | --- | --- |
| RNLS (Median) | 43.10% (20.16%-56.77%) | 27.78% (9.69%-53.48%) |
| RNLS (AUC) | 70.69% (57.27%-81.91%) | 66.67% (40.99%-86.66%) |
| CA19-9 | 0.00% (0.00%-45.93%) | 50.00% (24.65%-75.35%) |

**Figure 6.**
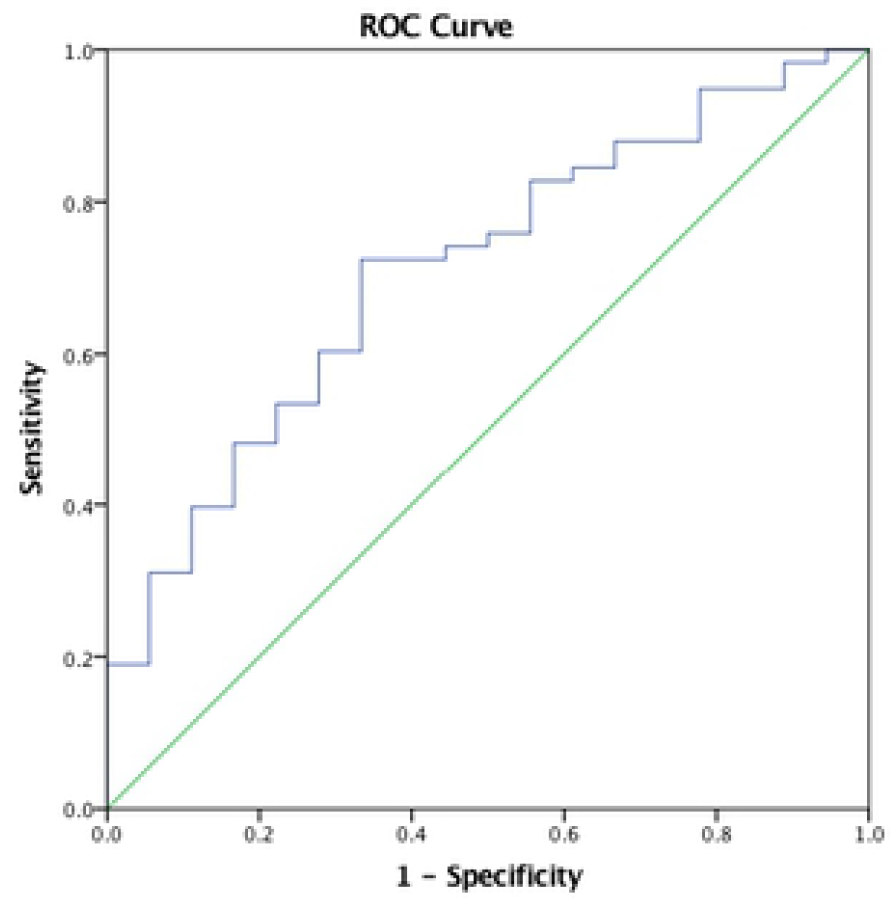
Plasma RNLS levels predict patients which patient will undergo resection in patients with locally advanced/borderline resectable pancreatic adenocarcinoma in ROC Curve. AUC (area under curve) = 0.674

## Discussion

Our data show that little to no RNLS immunoreactive protein was present in normal pancreatic tissue but that RNLS was detectable to varying degrees in premalignant PDAC and in PDAC tissues. The finding suggests a potential role for RNLS in the development of PDAC. Additionally, we found that high plasma RNLS levels corresponds to worse PDAC tumor characteristics, worse overall survival, and less resectability in a subset of patients with LA/BR PDAC.

In a subset of samples almost entirely represented by chronic pancreatitis, RNLS was also present in stromal cells, including inflammatory cells, fibroblasts, and stellate cells with a myofibroblastic phenotype. Stellate cells have important roles in the deposition of the extracellular matrix (stellate cells) and signaling to PDAC cells. The presence of RNLS in stellate cells in at least one chronic pancreatitis patient suggests that this protein could have a role in stellate cell function. Previous studies have shown that tumor microenvironment may exert selective pressures, including host immune response, proliferative or survival ability of cancer cells, or physiological restraints, which lead to the predominance of highly malignant PDAC cells(19, 20). We have previously shown that RNLS is present in tumor-associated macrophages adjacent to melanoma, which can promote tumor growth through a STAT3-mediated mechanism (21). Therefore, these findings suggest that RNLS could play a role in the tumor microenvironment during PDAC development. Future studies will determine the identity of the RNLS-labeled stromal cells that do not correspond to stellate cells.

Though essentially absent in other normal pancreatic tissues, RNLS labeling was present in the normal pancreas of patients who had undergone splenectomy with distal pancreatectomy for abdominal trauma. This difference likely reflects non-histologic pancreatic injury associated with abdominal trauma and non-specific increases in tissue RNLS as an acute phase reactant (22, 23).

A major finding of our study is that plasma levels of a select form of plasma RNLS correspond to PDAC prognosis. High plasma RNLS levels were also associated with worse overall survival, angiolymphatic invasion, node positive disease, including sentinel hepatic artery lymph node positive disease, and metastasis. Plasma RNLS levels were not associated with tumor size or surgical margin status among patients treated with Whipple or distal pancreatectomy surgery. It may be relevant that node positivity and metastasis reflect the clinically prognostic finding of tumor spread outside of the pancreas but that the effect of tumor size and margin status on prognosis in PDAC is controversial (24, 25). Overall, these findings suggest that plasma RNLS levels correlate with poor prognostic histopathologic features in PDAC.

Previous studies have suggested that tissue expression of RNLS is higher in select cancers than in benign tissues, including PDAC.(26-30) Tissue RNLS has been also described as a survival factor during ischemic or toxic injury and as a cytokine that activates several pathways, including PI3K/AKT, MAPK, p-ERK1/2, protein kinase B, and JAK/STAT pathways (28-30). In PDAC tissue studies, high RNLS tumor expression was associated with worse overall survival (8). Conditions that block RNLS can inhibit both in vitro and in vivo PDAC tumor growth (8). Further studies would benefit from understanding the mechanism of elevated plasma RNLS levels in patients with PDAC, whether different forms of RNLS have distinct tumor effects, whether tissue RNLS contributes the levels of plasma RNLS or if represents a host response to the tumor. Knowledge of whether plasma RNLS levels change with PDAC resection and can predict recurrence should also be clinically useful.

When considering patients who presented with LA/BC PDAC, our data show that reduced plasma RNLS, but not CA19-9, can predict both whether LA/BR PDAC patients will be subject to resection and their overall survival; CA-19 blood levels were not found to be predictive of these outcomes. Specifically, low plasma RNLS levels at diagnosis were associated with eventual eligibility for PDAC resection and with increased overall survival. There was a more than 2-fold increase in the rate of resection in patients with low plasma RNLS levels compared to high plasma RNLS levels. Previous studies have found survival improvements with resection in appropriately selected patients with locally advanced/borderline resectable PDAC (31). Though our study includes a limited number of patients, we found a non-significant trend towards improved survival in patients undergoing resection. Future studies should prospectively assess the role of RNLS in resection clinical decision making and survival outcomes. Together with tumor characteristic findings in resected PDAC samples, these findings suggest that plasma RNLS levels may reflect a tumor biology that predict outcome independent of radiologic tumor size, location, or vessel encasement at presentation. Plasma RNLS levels could also reflect characteristics that were not analyzed such as genetic, epigenetic or stromal microenvironment characteristics that portend worse outcomes.

Our study has a few limitations. First, the small sample size of each tissue type and immunohistochemical protocol limits statistically significant comparisons between each group the levels of RNLS in pancreatic tissue. Future studies using greater numbers of tissue samples and an automated immunohistochemistry platform are needed to better compare tissue levels of RNLS across tissues. For plasma RNLS studies, the retrospective cohort study design limited the ability to accurately assess survival over long periods of time to predict resectability without bias from independent treatment team decisions. We were also limited in analyzing factors that may play a role in determining a patient’s candidacy for surgery such as comorbidities, fitness, and anatomical limitations that were not uniformly present in chart review. Without serial plasma RNLS measurements or concurrent tissue expression of RNLS on plasma RNLS samples, we cannot correlate plasma RNLS with underlying PDAC tissue expression. The small sample size of this study also limited power. A larger sample size may have yielded a potential relationship between CA19-9 and RNLS that was not seen here among LA/BR patients. Additionally, for example, we found the median survival among patients who underwent resection versus no resection was 29.67 months vs. 30.57 months (p = 0.060; data not shown). This small sample size makes it difficult to evaluate this parameter in this cohort. Finally, there may have been selection bias associated with the type of patients who obtained care at our institution and subsequently were included in the biobank.

In conclusion, a key finding of our study is that RNLS is increased in both premalignant and malignant PDAC tissue compared to normal pancreas, suggesting a potential role for RNLS in the early development of PDAC. In addition, we found that the relationship between plasma RNLS levels and clinical outcomes of patients with PDAC which complements published data that correlated tissue RNLS expression and PDAC survival. These findings suggest that plasma RNLS could be used as a predictive biomarker in patients with PDAC and guide therapies such as resectability in LA/BR PDAC. The RNLS levels in tissue and plasma suggest a potential pathophysiological mechanism of RNLS for the development of PDAC and severe PDAC disease. Further studies should explore the potential mechanism of action of RNLS in pre-malignant pancreatic tissue and its expression in stromal cells including stellate cells.

Additionally, further studies should also explore disease progression patterns, plasma RNLS levels and tumor RNLS expression after resection, and the origin of plasma RNLS compared to tumor expression of RNLS. To further assess the ability of plasma RNLS to predict resectability in LA/BR pancreatic disease, larger prospective studies are needed that examine changes in plasma RNLS levels following neoadjuvant treatment. If plasma and tissue RNLS indeed reflects tumor biology and pathophysiology, it holds promise as a guide to surgical interventions and potential therapies that inhibit the pro-survival effects of RNLS in PDAC.

## ACKNOWLEDGEMENTS

We would like to acknowledge the staff members of the Yale Gastrointestinal Tumor Biorepository for providing access to patient samples for this study.

## FUNDING/FINANCIAL SUPPORT

This study was financially supported by the U.S. Department of Defense (DOD) Cancer Program 2019 (W81XWH1910439) to FG, GD, XG, and MR for salary support, laboratory analyses/equipment and materials necessary for this study and to MW for an NIH-NIDDK medical student fellowship award (DK007107). Funding from the Henry J. and Joan W Binder endowment also supported this work. The authors also wish to thank Christine Shugrue and Thomas Kolodecik for providing technical and constructive comments.

## PRIMARY DATA REPOSITORY

The original can be accessed at: DOI 10.17605/<u>OSF.IO/XXXX</u>

## COI/DISCLOSURES

G V Desir is a named inventor on several issued patents related to the discovery and therapeutic use of renalase. Renalase is licensed to Bessor Pharma, and G V Desir holds an equity position in Bessor and its subsidiary Personal Therapeutics.

